# Modulation of Giant Depolarizing Potentials (GDPs) in Human Large Basket Cells by Norepinephrine and Acetylcholine

**DOI:** 10.1101/2023.01.02.522475

**Authors:** Danqing Yang, Guanxiao Qi, Jonas Ort, Victoria Witzig, Aniella Bak, Daniel Delev, Henner Koch, Dirk Feldmeyer

**Affiliations:** Research Center Juelich, Institute of Neuroscience and Medicine 10, Research Center Juelich, 52425 Juelich, Germany; Department of Neurosurgery, Faculty of Medicine, RWTH Aachen University Hospital, Aachen, Germany; Department of Neurology, RWTH Aachen University Hospital, 52074 Aachen, Germany; Department of Neurology, Section Epileptology, RWTH Aachen University Hospital, 52074 Aachen, Germany; Department of Psychiatry, Psychotherapy, and Psychosomatics, RWTH Aachen University Hospital, 52074 Aachen, Germany; Jülich-Aachen Research Alliance, Translational Brain Medicine (JARA Brain), Aachen, Germany; Neurosurgical Artificial Intelligence Laboratory Aachen (NAILA), RWTH Aachen University Hospital, 52074 Aachen, Germany; Center for Integrated Oncology, Universities Aachen, Bonn, Cologne, Düsseldorf (CIO ABCD), Germany

**Author notes:** Correspondence should be addressed to Dirk Feldmeyer at or.

## Abstract

Rhythmic brain activity has been implicated in many brain functions and it is sensible to neuromodulation, but so far very few studies have investigated this activity on the cellular level *in vitro* in human tissue samples. In this study we revealed and characterized a novel rhythmic network activity in human neocortex. Intracellular patch-clamp recordings showed that giant depolarizing potentials (GDPs) were frequently found in human cortical neurons. GDPs appeared in a low frequency band (∼ 0.3 Hz) similar to that described for slow oscillations *in vivo* and displayed large amplitudes and long decay times. Under the same experimental conditions, no rhythmic activity was found in L2/3 of the rat neocortex. GDPs were predominantly observed in a subset of L2/3 interneurons considered to be large basket cells based on previously described morphological features. In addition, GDPs are highly sensitive to norepinephrine (NE) and acetylcholine (ACh), two neuromodulators known to modulate low frequency oscillations. NE increased the frequency of the GDPs by enhancing β-adrenergic receptor activity while ACh decreased GDP frequency through M_4_ muscarinic receptor-activation. Multi-electrode array (MEA) recordings demonstrated that NE promoted synchronous oscillatory network activity while the application of ACh led to a desynchronization of neuronal activity. Our data indicate that the human neocortex is more prone to generate slow wave activity, which was reflected by more pronounced GDPs in L2/3 large basket cells. The distinct modulation of GDPs and slow wave activity by NE and ACh exerts a specific modulatory control over the human neocortex.

## Introduction

Rhythmic brain activity has been implicated in many brain functions, from sensory processing to memory consolidation [1-4]. Sleep states, in particular, are largely dependent on synchronized neuronal activity generated within the cerebral cortex. In the neocortex, non-rapid eye movement (NREM) sleep is dominated by slow oscillations (< 1 Hz). The intracellular correlates of these slow oscillations - termed ‘Up’ and ‘Down’ states - represent periods of activity interspersed with periods of relative silence [5]. In contrast, during the rapid-eye-movement (REM) sleep and active wakefulness, the neocortex shows persistent but asynchronous activity of a low amplitude and a high frequency [6, 7]. However, slow oscillations are observed not only during physiological conditions like NREM but also during anesthesia or in pathological conditions such as stroke or traumatic brain injury [8-10]. Understanding the dynamics of slow oscillations in cortical networks and how they vary during transitions between different behavioral states is crucial to decipher how these networks function.

Norepinephrine (NE) and acetylcholine (ACh) are two well-known neuromodulators that have long been implicated in altering network dynamics during different behavioral states [11-13]. By stimulation of brainstem noradrenergic and cholinergic nuclei, NE and ACh release is known to modulate oscillations and sleep substates [14-16]. Cholinergic basal forebrain and brainstem neurons show increased firing during desynchronization of rhythmic activity, in turn promoting wakefulness and attention [17]. On the other hand, noradrenergic neurons were proved to be highly active and show coordinated activity variation during NREMs [16, 18]. Activation of noradrenergic neurons in the locus coeruleus (LC) contributes to transitions between ‘Down’ and ‘Up’ states [19]. With only a limited number of studies available for human neurons [20, 21], our current knowledge about the neuromodulation of the neocortex by NE and ACh is derived from in vitro/in vivo experiments in rodents. However, the density of neuromodulatory afferents is much higher in the neocortex of primates than rodents [22, 23]. Therefore, it is essential to investigate the neuromodulation of network activity in human neocortical tissue to reveal their underlying mechanisms.

Here, we report that in acute human brain slices, spontaneous rhythmic giant depolarizing potentials (GDPs) were detected in neocortical layer 2/3 in the absence of pharmacological manipulation or external stimulation. These GDPs are more prevalent in human interneurons than pyramidal cells. Of note, GDPs were not observed in rat brain slices under identical conditions. Human interneurons showing GDPs are basket cells with large dendritic and axonal arborization domains, hence displaying distinct morphologies when compared to interneurons without GDPs. Moreover, GDPs are highly sensitive to noradrenergic and cholinergic modulation with NE enhancing and ACh reducing GDP frequency. MEA recordings revealed a clear enhancement of spiking synchronization by NE while ACh desynchronized neuronal firing. Taken together with the dynamics of GDPs at the network level as well as their responses to neuromodulators, we consider GDPs as ‘Up’ states during slow oscillations maintained in vitro. We demonstrated that as a hallmark of brain activity, slow oscillations are under antagonistic neuromodulatory control of NE and ACh in human brain.

## Results

### Giant depolarizing potentials (GDPs) are observed in a subset of human cortical L2/3 neurons

Acute brain slices were prepared from tissue blocks following brain surgery (resection of epileptic and tumor foci) in 21 patients aged 8-75 years (**Tab. 1**). Blocks of neocortical tissue were removed during the surgical approach to the pathological brain region. All tissue used here was sufficiently distant from the tumor and/or the epileptogenic focus, thus consisting of normal cortex. Brain tissue was surgically removed from frontal or temporal cortex, however, in one case from the occipital and one from the parietal cortex (**Tab. 1**). Whole-cell current clamp recordings with simultaneous biocytin fillings were performed from human L2/3 cortical neurons allowing post hoc identification of their morphologies. 60 pyramidal cells (PCs) and 52 interneurons (INs) were identified by their electrophysiological properties and morphological features (**Fig. 1e**). In a subset of human L2/3 neurons (20 out of 120 neurons), giant depolarizing potentials (GDPs) were observed against the background of normal excitatory postsynaptic potentials (EPSPs). These large, complex events occurred spontaneously and rhythmically; in some neurons, they directly triggered action potential (AP) discharges when the depolarization reached the AP threshold (**Fig. 1a**). GDPs were not associated with a specific brain region, gender, age or pathophysiology (**Tab. 1**). For each neuron exhibiting GDPs, we analyzed the excitatory postsynaptic activity (continuous current-clamp recording, 100 s). GDPs and normal EPSPs were analyzed separately. In L2/3 interneurons, GDPs exhibited an average amplitude of 10.2 ± 3.1 mV (n = 448 events in 14 neurons), which was significantly larger than that of unitary EPSPs (2.1 ± 1.1 mV, n = 14154 events in 14 neurons). Dramatic differences in dynamic properties were also observed between GDPs and normal EPSPs. The decay time was significantly longer for GDPs (154.0 ± 44.7 ms) than for spontaneous EPSPs (41.2 ± 14.7 ms) (**Fig. 1b, d**). GDPs occurred at a low frequency ranging between 0.11 and 0.70 Hz (mean = 0.32 ± 0.18 Hz). In contrast, normal EPSPs had a much higher frequency of 10.11 ± 6.14 Hz (**Fig. 1c-d**) suggesting that the functional mechanisms underlying GDP generation are fundamentally different from those of normal EPSPs. Of note, GDPs were more frequently observed in interneurons than PCs (15 out of 52, 28.8% vs. 5 out of 68, 7.4%, respectively; **Fig. 1f**). GDP frequency or rise time in human L2/3 PCs and interneurons was similar, however, a decreased GDP amplitude and a prolonged decay time were observed for PCs (**Supplementary Fig. S1**). We found that GDPs were detected only within the first 6 hours after slicing but not thereafter (see also **Supplementary Fig. S2**).

**Fig. 1.**
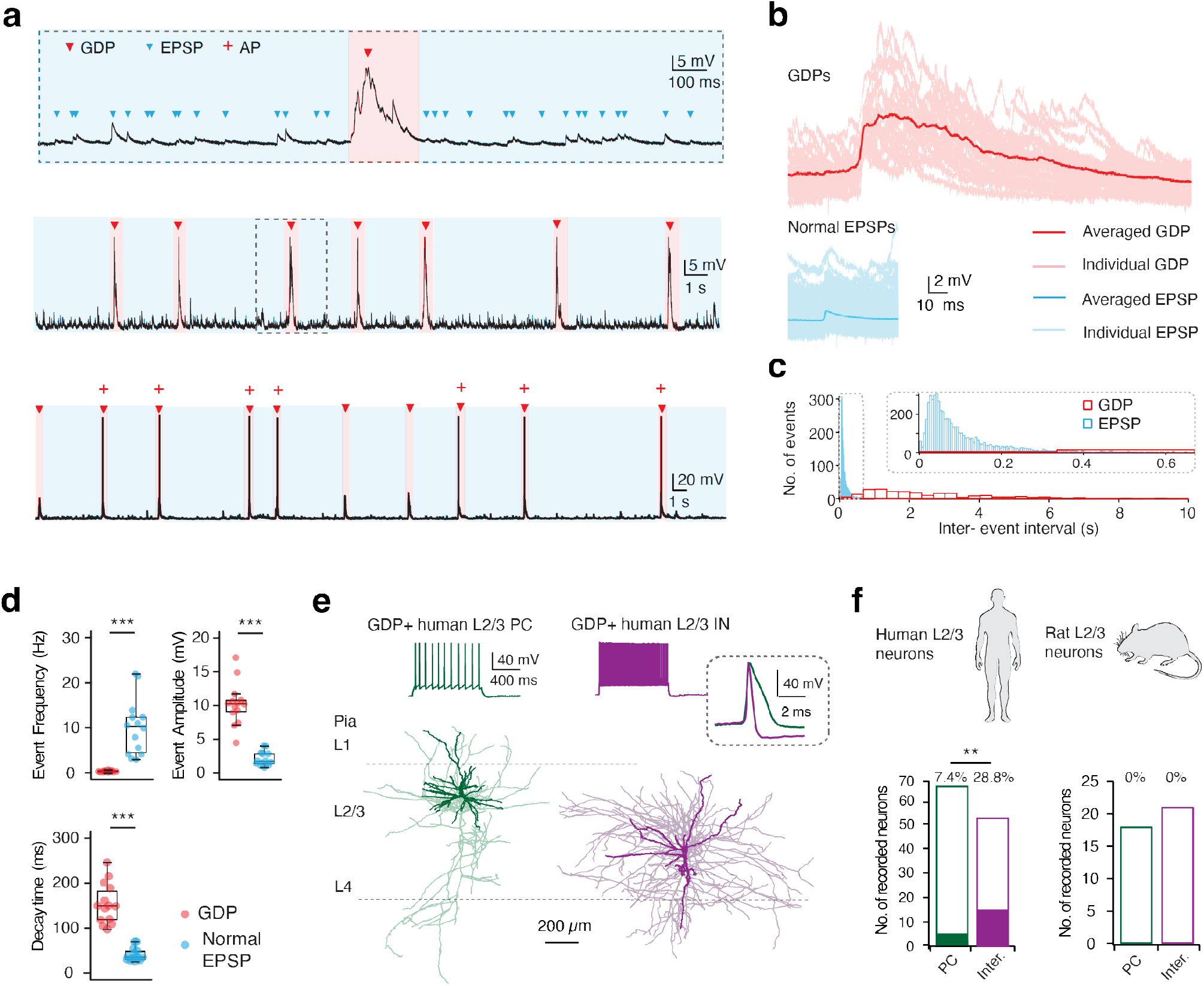
GDPs are identified in a subset of human neocortical L2/3 neurons. **a** Whole-cell intracellular recordings from a L2/3 interneuron in the human neocortex. Top trace: a typical GDP in the temporal resolution of 2 s. The same GDP is shown in the middle trace on an expanded time scale. Normal excitatory postsynaptic potentials (EPSPs) are marked by blue while GDPs are marked by red arrowheads. Middle trace: continuous 30 s recording of spontaneously, rhythmically occurring GDPs. Bottom trace: spontaneous GDPs triggering action potential discharge. **b** Top: Mean and individual GDPs (n = 29) are superimposed and given in dark and light red, respectively. Bottom: Mean and individual EPSPs (n = 559) are superimposed and given in dark and light blue, respectively. Events were extracted and analyzed from 100 s continuous recordings. **c** A 300 s recording was obtained from the same neuron in (a) and (b) and interval histograms of events are shown. The histograms of GDPs and normal EPSPs were constructed with 33 ms and 5 ms bins, respectively. Inset, EPSP histogram at an expanded time scale. **d** Box plots comparing event frequency, amplitude and decay time for GDPs and normal EPSPs. n = 14 neurons for each group; *** P < 0.001 for the Wilcoxon Mann–Whitney U test. **e** Top: Representative firing patterns of a human L2/3 PC and an interneuron exhibiting spontaneous GDPs. The inset shows the first AP elicited by rheobase current at high temporal resolution. Bottom: Corresponding morphological reconstructions of the neurons shown above. The somatodendritic domain is given in a darker, the axons in a lighter shade.

To investigate whether GDPs are neuronal network events specific to human brain slices, recordings were performed from L2/3 neurons in acute brain slices of adult rat prefrontal and temporal cortex. No GDPs were detected in any rat L2/3 PCs (n = 18, **Fig. 1f** & **Supplementary Fig. S1**). Although we observed sporadic GDP-like events in 2 out of 21 L2/3 interneurons, these events showed no rhythmicity or persistency and therefore were markedly different from typical GDPs in human neurons (**Supplementary Fig. S1**). Taken together, our data suggests that GDPs are giant, rhythmic postsynaptic depolarizing events unique to human neocortex that occur more frequently in interneurons than in PCs.

It has been reported that slow rhythmic activity is initiated and prominent in infra-granular layers and then propagates to supra-granular layers [24]. To investigate the network activity in all the neocortical layers, extracellular recordings were performed using the 256–multi-electrode array (MEA) on human cortical brain slice cultures. Local field potentials (LFPs) were detected both in L2/3 and L5/6 (**Supplementary Fig. S3a-b**). Therefore, whole-cell recordings were performed from human L6 cortical neurons and typical GDP activity was observed in L6 interneurons as well (**Supplementary Fig. S3c-d**). Taken together, GDPs are not confined to L2/3 but appear across different neocortical layers.

### GDPs are complex network events depending on presynaptic glutamatergic release

GDPs are complex events, often comprising several depolarizing components. Previous studies suggested that human L2/3 PCs form stronger, more reliable connections compared to rodents [25, 26]. Such strong connections make a direct postsynaptic AP more likely and could contribute to synchronized AP firing in the local neuronal microcircuitry. Unlike spontaneous unitary synaptic inputs triggered by individual presynaptic APs, we hypothesize that GDPs are induced by near-synchronous firing in presynaptic glutamatergic neurons (**Fig. 2a**). As an initial test of this hypothesis, we performed a time-frequency analysis of the spontaneous synaptic activity in GDP-positive interneurons. Low-frequency GDPs and high-frequency EPSPs can be clearly discriminated (**Fig. 2b**). The onset of a GDP was always accompanied by a superposition of small amplitude events, indicating that GDPs are network-driven synchronous oscillations (**Fig. 2b**). To study the presynaptic mechanisms underlying GDP generation, either 0.5 µM TTX or 10 µM CNQX was bath-applied. Both TTX and CNQX completely abolished GDPs, suggesting that GDPs are AP-dependent synaptic events relying on presynaptic glutamatergic release (**Fig. 2c**). Giant depolarizing events were reported previously for the rat hippocampus and neocortex during the first postnatal week, and were elicited by γ-aminobutyric acid (GABA) which is a depolarizing transmitter at early developmental stages [27-30]. In our experiments bath application of GABA (1 µM) did not affect the GDP frequency, indicating that these events have a different underlying mechanism than those found in immature rat hippocampus.

**Fig. 2.**
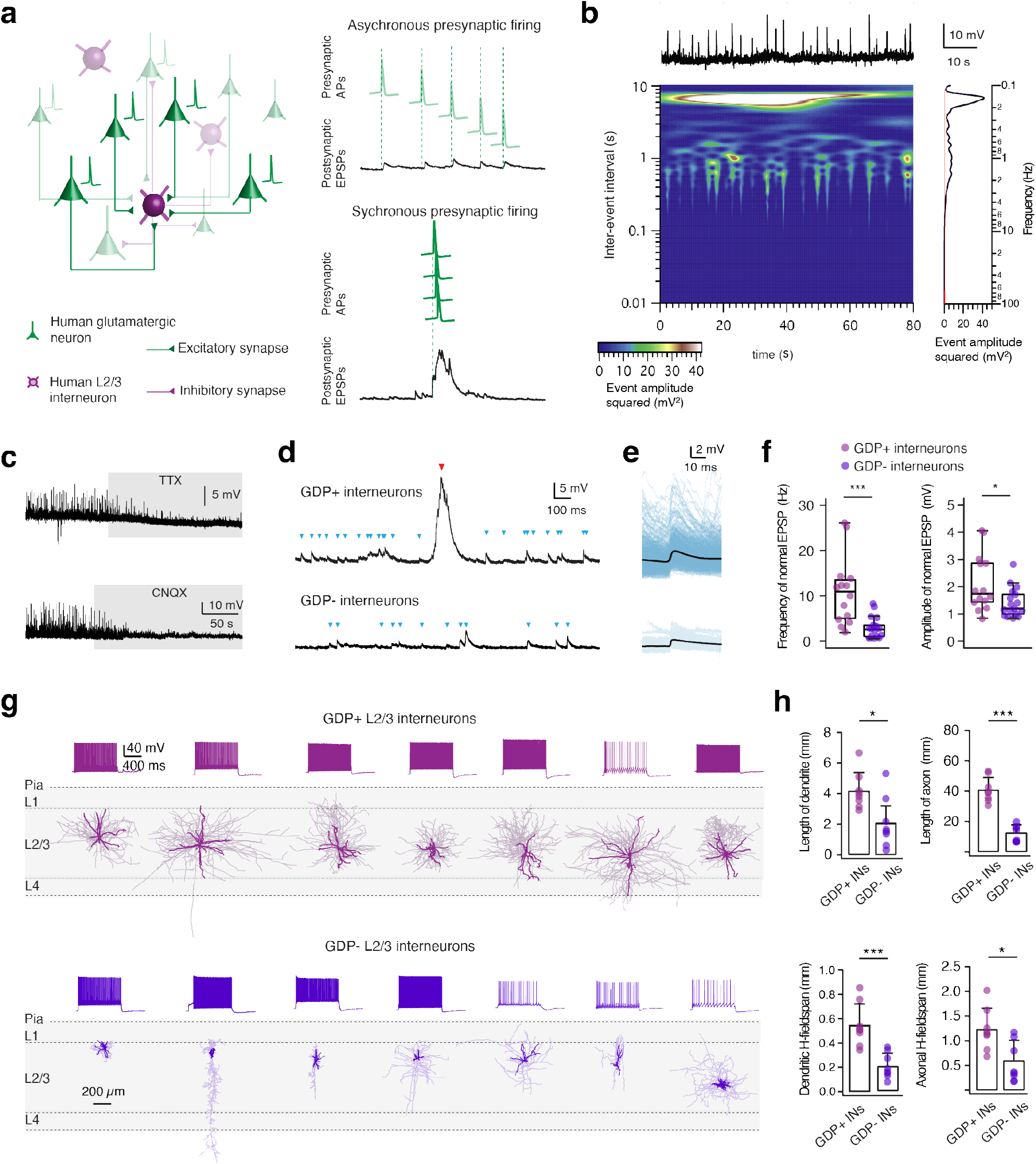
GDPs are complex network events depending on glutamatergic transmission and human L2/3 interneurons showing GDPs are large basket cells. **a** Left: Synaptic wiring scheme between human L2/3 interneurons and cortical PCs. Right: Diagram summarizing the possible mechanism of the generation of normal EPSPs and a GDP. Presynaptic asynchronous APs are shown in light green while synchronous APs are in dark green. **b** Time-frequency representation of excitatory spontaneous activities in a GDP-positive human L2/3 interneuron. Top plot: Original current clamp recordings in time from a human L2/3 interneuron. Central plot: A time-frequency decomposition of the recording shown above. Squared event amplitude is depicted by heat map colors. Right plot: Amplitude spectrum of excitatory spontaneous activities. The spectrum is shown vertically in frequency (0.05-100 Hz), the red line **c** Representative current clamp recordings showing block of GDPs in human L2/3 interneurons by TTX (0.5 µM, top trace) and CNQX (10 µM, bottom trace). **d** Representative continuous recordings from GDP-positive (top trace) and GDP-negative (bottom trace) human L2/3 interneurons. Normal EPSPs are marked in blue while GDPs are marked in red. **e** Normal EPSPs were extracted and analyzed from the same GDP-positive interneuron (top) and GDP-negative interneuron (bottom) in d. The average and individual EPSP traces are superimposed and given in black and blue, respectively. **f** Box plots comparing the frequency and amplitude of normal EPSPs for GDP-positive and GDP-interneurons. n = 14 for GDP-positive interneurons and n = 17 for GDP-negative interneurons. *P < 0.05, *** P < 0.001 for the Wilcoxon Mann–Whitney U test. **g** Representative morphological reconstructions and the corresponding firing patterns of seven GDP-positive (top) and seven GDP-negative (bottom) interneurons. The somatodendritic domain is shown in black and axons are shown in gray. **h** Histograms comparing several morphological properties of GDP-positive and GDP-negative human L2/3 interneurons. n = 8 for each group. *P < 0.05, *** P < 0.001 for the Wilcoxon Mann– Whitney U test.

Since GDP positive neurons could represent a group of highly connected cells in the neuronal network, we examined the timing and magnitude of EPSPs between GDP-positive and GDP-negative L2/3 interneurons. Interestingly, GDP-positive interneurons showed a much higher frequency of EPSPs compared to GDP-negative interneurons (11.0 ± 7.7 vs. 2.9 ± 2.4 Hz, P < 0.001). In addition, we revealed a normal EPSP amplitude which was on average 1.5 times larger in GDP-positive interneurons compared to GDP-negative interneurons (GDP + INs: 2.1 ± 0.9 mV, GDP - INs: 1.4 ± 0.5 mV, P < 0.05) (**Fig. 2d-f**). Overall, our data demonstrates that GDP-positive interneurons receive strong excitatory input and are actively involved in neuronal network. This lends further support to the idea that human neocortex is more prone to generate GDPs than rodent neocortex because of the remarkable strong excitatory-to-inhibitory connectivity in human L2/3 neocortex [26, 31].

### Interneurons showing GDPs are large basket cells

Neocortical interneurons can be divided into several distinct subtypes based on their electrophysiological, morphological and transcriptional properties [32-34]. To investigate whether GDPs were observed in a defined interneuron subtype, we analyzed the electrophysiological properties and performed 3D morphological reconstructions of L2/3 interneurons with and without GDPs. There is no clear correlation between GDP occurrence and neuronal firing pattern. Both GDP-positive and -negative interneurons exhibit diverse firing patterns comprising typical fast-spiking (FS) and non-fast spiking (nFS, which includes adapting, late spiking, etc.) firing behavior as described in the literature (**Fig. 2g**; **Supplementary Tab. S1**) [35, 36]. However, after detailed analysis of passive and active properties, GDP-positive interneurons were found to exhibit a smaller input resistance when compared to GDP-negative interneurons (192.4 ± 85.1 vs. 298.5 ± 117.9 MΩ, P < 0.05). We next studied the morphologies of GDP-positive and GDP-negative L2/3 interneurons. Notably, GDP-positive interneurons were mostly large basket cells with dense and broad dendritic and axonal domains. In contrast, GDP-negative interneurons comprised multiple morphological subtypes including small basket, bipolar, double bouquet and neurogliaform cells (**Fig. 2g**). GDP-positive interneurons showed significantly longer dendrites and axons than GDP-negative interneurons (4.2 ± 1.2 vs. 2.1 ± 1.1 mm for dendritic length, P < 0.05; 40.8 ± 8.2 vs. 12.5 ± 5.4 mm for axonal length, P < 0.001). The horizontal dendritic and axonal fieldspan was wider for GDP-positive than for GDP-negative interneurons (546 ± 176 µm vs. 212 ± 104 µm for dendrites and 1228 ± 427 µm vs. 604 ± 405 µm for axons, respectively; **Fig. 2h**). Moreover, a larger vertical dendritic fieldspan was also observed for GDP-positive neurons (652 ± 194 vs. 308 ± 217 µm, P < 0.01). More details regarding morphological properties and statistical comparisons are given in Supplementary Table S1.

Parvalbumin (PV)-expressing GABAergic interneurons are known to provide perisomatic inhibition onto PCs and contribute to cortical network oscillations [37]. A large portion of basket cells in the human neocortex are PV-expressing neurons displaying a FS firing pattern [38, 39]. To identify the expression of PV in human L2/3 interneurons, we performed whole-cell recordings with simultaneous filling of biocytin and the biocytin-conjugated fluorescent Alexa Fluor 594 dye. Subsequently, brain slices were processed for PV immunofluorescence staining. We found that FS interneurons showing GDPs were PV-positive, while the nFS interneurons showing GDPs were PV-negative (**Supplementary Fig. S4**). This indicates that, although GDP-positive interneurons show uniform morphologies, they contain more than one transcriptional type of basket cells including FS PV-positive cells and possibly also nFS cholecystokinin (CCK)-positive interneurons [40, 41].

### Norepinephrine (NE) induces GDPs or enhances their frequency via β-adrenergic receptors

In a subset of human L2/3 interneurons (n = 7) GDPs were induced following bath application of 30 µM NE (**Supplementary Fig. S5**). The morphological and functional properties of these neurons suggest that they are large basket cells exhibiting high-frequency background EPSPs, similar to those L2/3 interneurons with spontaneous GDPs under control conditions. Thus, they were included in the statistical analysis of electrophysiological and morphological properties of GDP-positive interneurons in **Fig. 2** and **Supplementary Tab. S1**. Furthermore, in L2/3 interneurons with spontaneous GDPs, 30 µM NE significantly increased GDP frequency. The NE-induced changes of GDP frequency were reversible following washout (**Fig. 3a**). A similar increase of the GDP frequency was also observed in the presence of only 10 µM NE (**Supplementary Fig. S6**). To examine whether NE specifically altered the neuronal excitability in GDP-positive interneurons, we measured NE-induced changes in the resting membrane potential (V_m_) but found no significant effect in both GDP-positive and GDP-negative interneurons (**Fig. 3b**).

**Fig. 3.**
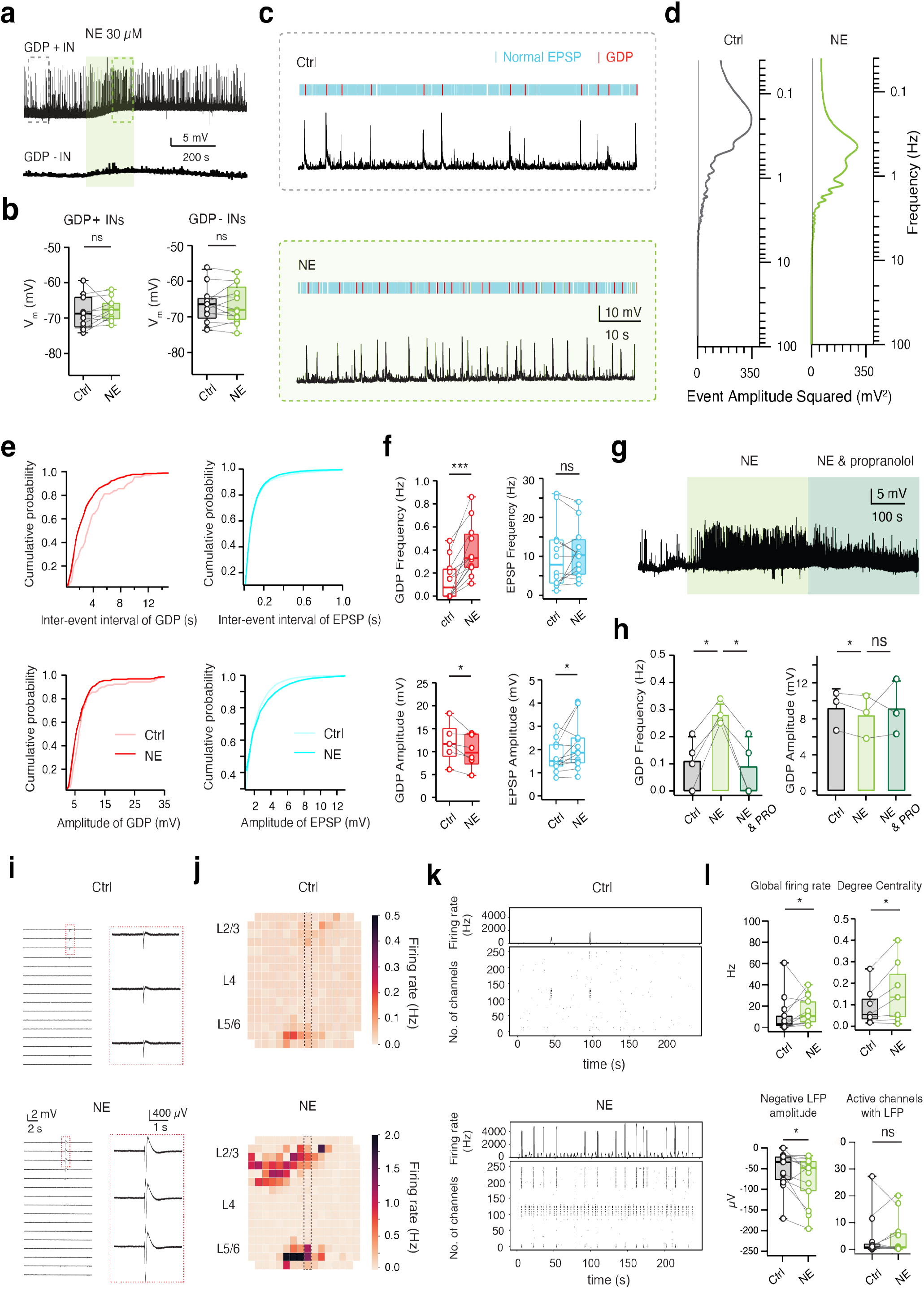
NE increases GDP frequency via activating β-adrenergic receptors. **a** Representative GDP-positive (GDP+) interneuron (IN; top trace) and GDP-negative (GDP-) interneuron (bottom trace) showing depolarizing responses following the bath application of 30 µM NE. The area in the dashed boxes is enlarged in c. **b** Summary plots showing the resting membrane potential (V_m_) of human L2/3 interneurons under control and in the presence of NE (n = 10 for GDP-positive interneurons and n = 12 for GDP-negative interneurons). ns (not significant); Wilcoxon signed-rank test. **c** 80 s recording from the same GDP-positive interneuron under control (top) and in the presence of NE (bottom). Normal excitatory postsynaptic potentials (EPSPs) are marked in blue while GDPs are marked in red. **d** Amplitude spectrum of excitatory spontaneous activities analyzed from recording traces in c. The spectrums are shown vertically in frequency (0.05–100 Hz) and the black lines represent baseline noise. **e** Cumulative distributions of inter-event intervals and amplitudes of excitatory spontaneous activity recorded in GDP-positive human L2/3 interneurons under control and NE conditions. **f** Box plots summarizing the NE effect on frequency and amplitude of GDPs and normal EPSPs in GDP-positive L2/3 human interneurons (n = 12). ns (not significant), *P < 0.05, ***P < 0.001; Wilcoxon signed-rank test. **g** Representative current-clamp recordings, following the bath application of NE, showing an increase of GDP frequency in a human L2/3 interneuron. The effect is blocked by the β-adrenoreceptor antagonist propranolol (20µM). **h** Summary histograms of GDP frequency and amplitude under control, NE and propranolol conditions. ns (not significant), *P < 0.05 for paired student t-test. **i** Representative voltage traces from 16 channels of the MEA under control conditions (top) and 30 µM NE (bottom) showing an increase in LFP amplitude. Red insets show an enlarged view of the same three electrodes in both conditions. **j** Heatmap of the average firing rate over the MEA grid from a five-minute recording in control condition and in the presence of NE showing an increase synchronous firing in L2/3 and deep layers. **k** Raster plots of the detected APs over a five-minute recording period under control condition and in the presence of NE showing an increase in synchronous firing.

GDPs and normal background EPSPs were analyzed separately for control conditions and in the presence of 30 µM NE (**Fig. 3c**). Our results show that NE differentially modulates GDPs and normal EPSPs in GDP-positive human L2/3 interneurons. Spectral analysis of the spontaneous activity in GDP-positive interneurons revealed that NE significantly increased the frequency of the GDPs (from 0.13 ± 0.17 Hz to 0.39 ± 0.23 Hz, n = 12 neurons, P < 0.001) without affecting the frequency of the EPSPs (10.16 ± 8.53 vs. 10.86 ± 6.52 Hz, P = 0.5796, **Fig. 3d**). In addition, NE slightly decreased the amplitude of GDPs (from 11.92 ± 4.07 to 10.02 ± 3.53 mV, P < 0.05) but increased that of normal EPSPs (from 1.69 ± 0.65 to 2.11 ± 1.02 mV, P < 0.05) (**Fig. 3e-f**). To determine the specific adrenergic receptor type which mediates the NE-induced increase in GDP frequency, either 2 µM prazosin (a selective antagonist of α_1_-adrenergic receptors) or 20 µM propranolol (a selective antagonist of β-adrenergic receptors) was co-applied with 30 µM NE following bath application of NE alone. While prazosin had no effect on the adrenergic response, propranolol completely blocked the NE effect on GDP frequency. The GDP frequency increased from 0.11 ± 0.09 to 0.28 ± 0.04 Hz during NE application and returned to control level (0.09 ± 0.10 Hz) following co-application of NE and propranolol (**Fig. 3g-h**). These experiments indicate that the NE-induced enhancement of the GDP frequency is mediated by activation of β-adrenoreceptors.

To determine the effect of NE on global cortical network activity, we recorded human cortical brain slice cultures on muti-electrode array (MEA) before and following bath application of NE and ACh. Under control conditions, all slices included in the analysis (n = 25, 6-16 days in vitro (DIV)) showed spontaneous network activity (**Fig.3 i-j** and **4 i-j**) with a mean firing rate over all channels of 10.5 ± 15.4 Hz, with 23.4 ± 42.7 (of 256) active channels and 5.7 ± 8.6 channels with detected local field potentials (LFPs; see methods for detail). As a summed signal from synchronous neuronal activity near the recording site, LFP reflects the dynamic flow of network activity. The mean amplitude of LFP was -57.8 ± 46.7 over a time period of 6-10 minutes. Bath application of NE (30 µM, n = 13) led to a significant increase in the global firing rate (from 10.5 ± 17.2 Hz to 14.8 ± 12.8 Hz, *p < 0,05) combined with a significant increase in the negative amplitude of the detected LFPs (from -53.2 ± 45.5 µV to -76.3 ± 59.4 µV, *p < 0.05) (**Fig. 3j-l**). To quantify the synchronicity of the neuronal network, we performed graph analytics. We calculated the degree of centrality for channels with simultaneous spiking activity as surrogate measure for the synchronicity. The mean degree centrality (MDC) for slices treated with NE increased from 0.09 ± 0.09 to 0.16 ± 0.15 (*p < 0.05, **Fig. 3l** and **Supplementary Fig. S7e**). The increase in activity was observed mainly with electrodes that were already active under control conditions or in close proximity to those (**Fig 3j**); no significant difference in the number of active channels with a LFP was detected (**Fig. 3l**). The increase in firing rate following bath application of NE was statistically not different for electrodes located either in L2/3 or L5/6.

### Acetylcholine (ACh) suppresses GDPs via the activation of M_4_Rs

ACh shifts cortical dynamics from a synchronous to asynchronous state and improves the signal-to-noise ratio of sensory signaling [42, 43]. To elucidate cholinergic effects on human L2/3 interneurons, 30 µM ACh was bath applied and ACh-induced changes in V_m_ were compared between interneurons with and without GDPs. ACh persistently depolarized all tested human L2/3 interneurons and was reversible following washout with ACSF. GDP-positive and GDP-negative L2/3 interneurons were depolarized by 4.0 ± 3.5 mV (n = 10) and 3.1 ± 2.6 mV (n = 12), respectively (**Fig. 4b**). No significant differences were detected in depolarization amplitude.

**Fig. 4.**
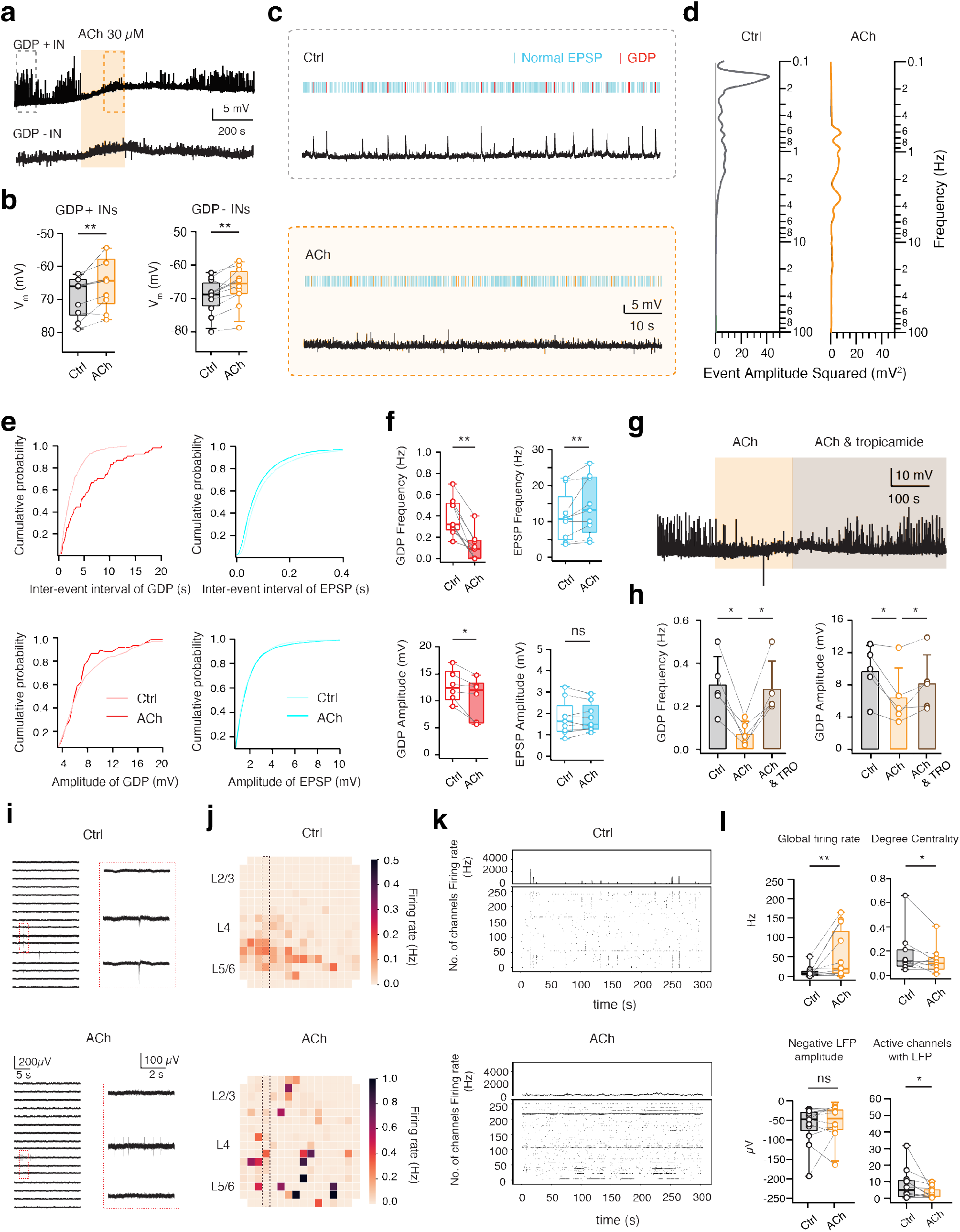
ACh suppresses GDPs via activating M_4_Rs. **a** Representative GDP-positive interneuron (top trace) and GDP-negative interneuron (bottom trace) show depolarizing responses following the the bath application of 30 µM ACh. The area in the dashed boxes is enlarged in c. **b** Summary plots showing the resting membrane potential (V_m_) under control conditions and in the presence of ACh in human L2/3 interneurons. n = 10 for each group. **P < 0.01; Wilcoxon signed-rank test. **c** A 80 s recording from a GDP-positive interneuron under control (top) and ACh conditions (bottom), respectively. Spontaneous EPSPs are marked in blue while GDPs are marked in red. **d** Amplitude spectrum of excitatory spontaneous activity analyzed from recording traces in c. Spectra are shown vertically in frequency (0.05–100 Hz), black lines represent baseline noise. **e** Cumulative distributions of inter-event interval and amplitude of spontaneous excitatory activity in GDP-positive human L2/3 interneurons under control conditions and in the presence of ACh. **f** Box plots summarizing the ACh effect on frequency and amplitude of GDPs and normal EPSPs in GDP-positive L2/3 human interneurons (n = 10). ns (not significant), *P < 0.05, **P < 0.01; Wilcoxon signed-rank test. **g** Representative current-clamp recordings with bath application of ACh showing a decrease of GDP frequency in a human L2/3 interneuron. The effect is reversed by 1 µM tropicamide. **h** Summary histograms of GDP frequency and amplitude under control conditions, in the presence ACh alone and together with the M_4_ mAChR antagonist tropicamide. *P < 0.05; Wilcoxon signed-rank test. **i** Representative voltage traces from 16 channels of the MEA under control conditions (top) and 30 µM ACh (bottom) showing block of LFPs by ACh. Red inserts show a detailed view of the same three electrodes in both conditions; note the switch from a synchronized firing pattern under control condition to a desynchronized pattern in the presence of ACh. **j** Heatmap of the average firing rate over the MEA grid from a five-minute recording in control condition and in the presence of ACh showing a decrease in temporal and spatial correlation of global AP firing. **k** Raster plots of the detected action potentials over the five-minute recording period under control condition and in the presence of ACh showing a decrease in synchronous firing. **l** Box plots displaying the group effects of ACh compared to control on the global firing rate (n =

We found that ACh application resulted in a marked suppression of both spontaneous (**Fig. 4a**) and NE-induced GDPs (**Supplementary Fig. S5**). To systematically study cholinergic modulation of spontaneous activity, we analyzed GDPs and background spontaneous EPSPs from continuous current-clamp recordings during 100 s epochs in each GDP-positive interneuron under control conditions and in the presence of ACh (**Fig. 4c**). Spectral analysis of spontaneous activity in GDP-positive interneurons indicates that ACh blocked large amplitude events but increased the frequency of small amplitude events (**Fig. 4d**). In fact, ACh either decreased (n = 6) or completely blocked (n = 4) the occurrence of GDPs, resulting in a reduction of GDP frequency from 0.40 ± 0.17 Hz to 0.13 ± 0.13 Hz. Of note, ACh at a lower concentration of 10 µM similarly reduced GDP frequency (**Supplementary Fig. S6**). In contrast, 30 µM ACh significantly increased the frequency of normal EPSPs from 11.18 ± 6.74 Hz to 14.12 ± 7.99 Hz (**Fig. 4e-f**). Our data suggests that ACh modulates unitary synaptic and GDP activity in a distinctive fashion, leading to a desynchronization of synaptic potentials thereby reducing GDP frequency or abolishing GDP occurrence altogether. Additionally, ACh slightly decreased GDP amplitude from 12.74 ± 2.94 mV to 10.37 ± 4.24 mV but had no effect on the amplitude of normal EPSPs (1.78 ± 0.80 vs. 1.78 ± 0.65 mV, P = 0.9856) (**Fig. 4e-f**). It has been reported that by blocking K^+^ conductances, ACh terminates ‘Up’ states via muscarinic receptors [14, 44]. Here we found that the cholinergic modulation of GDPs was blocked by administration of a muscarinic M_4_ receptor (M_4_R) antagonist, tropicamide, but not by mecamylamine, a general nicotinic antagonist. The GDP frequency, which was reduced by ACh, increased to control level in the presence of 1 µM tropicamide (from 0.07 ± 0.06 to 0.28 ± 0.13, P < 0.05). Likewise, the ACh-induced enhancement of the GDP amplitude was reversed as well (from 6.44 ± 3.65 to 8.16 ± 3.54 mV, P < 0.05, **Fig. 4g-h**). These results suggest that ACh suppresses GDPs mainly via the activation of M_4_Rs.

To further test how ACh modulates cortical network activity, we measured the effect in human cortical slice cultures using a 256 MEA recording system. Bath application of ACh (15-30 µM, n = 12) led to a significant increase in the mean AP firing rate (from 9.7 ± 13.7 Hz to 50.1 ± 62.2 Hz, **p < 0.01, **Fig. 4l**). However, the increase in firing rate was patchy and disjunct (**Fig. 4j**) and the temporal and spatial correlation observed under control conditions was not preserved in the presence of ACh (**Fig. 4j-k**). Furthermore, the enhanced firing rate was not accompanied by an increase of the LFP amplitude (underlying synchronous network events), but with a significant decrease of the number of active electrodes showing LFPs (**Fig. 4l**). Network activity became asynchronous as the decrease in the degree of centrality from 0.17 ± 0.18 to 0.12 ± 0.11 (*p < 0.05) after application of ACh indicates (**Fig. 4l and Supplementary Fig. S7f**). No significant difference was found between the increase in firing rate between electrodes located in L2/3 or L5/6 in response to the bath application of ACh.

To further probe how AP firing was modulated by NE and ACh, 30 µM NE and 30 µM ACh were applied on L2/3 interneurons showing GDP-induced AP firing. We found that NE increased the GDP frequency and in turn enhanced the GDP-induced AP firing rate. In contrast, application of ACh strongly depolarized neurons and thereby boosted the general AP firing rate in a random fashion. AP firing was, however, no longer correlated, thereby preventing the generation of GDPs (**Supplementary Fig. S8**). This indicates that NE and ACh enhance AP firing rates in L2/3 interneurons through distinct mechanisms.

## Discussion

In this study, we uncovered and characterized rhythmic network events, revealing cell type-specific network activity in layer 2/3 of human neocortex, here coined Giant Depolarizing Potentials (GDPs). GDPs appeared in a low frequency range (0.1-0.7 Hz) corresponding to the frequency band for slow oscillations, which is a hallmark of NREM sleep and anesthesia in human brain [45, 46]. GDPs displayed large amplitudes, long decay times and sometimes triggered AP firing. Although L2/3 PCs occasionally showed GDPs, they were predominantly observed in a subset of L2/3 interneurons exhibiting dendritic and axonal morphologies similar to those of large basket cells [47]. In contrast to human neocortex, GDPs were not observed in L2/3 of rat frontal or temporal cortex in our experiments, suggesting a species specificity. Our data indicates that GDPs are evoked by presynaptic AP firing in glutamatergic neurons. In addition, the synchronous network events and asynchronous unitary synaptic inputs were differentially modulated by NE and ACh. NE increased GDP frequency via activating β-adrenergic receptors without affecting the frequency of normal EPSPs. Conversely, ACh decreased GDP frequency by M_4_ muscarinic receptor activation but enhanced the frequency of normal EPSPs. Data obtained from MEA recordings further demonstrated that NE boosted near-synchronous AP firing while ACh desynchronized network activity. The distinct modulation of such activity by NE and ACh implies specific modulatory mechanisms in the human neocortex, shedding light on mechanisms of synchronized neuronal activity in human neocortex which is associated with different behavioral states.

GDPs have been observed in the immature rodent hippocampus and neocortex as network-driven synaptic events. Their initiation requires excitatory GABAergic transmission which promotes voltage-dependent AP bursts in immature pyramidal neurons [30, 48]. The GDPs discovered and characterized in the present work were recorded in L2/3 of human neocortex and are dependent on glutamatergic transmission and are not affected by administration of GABA, suggesting a distinct mechanism in adult human neocortex. In acute human, monkey, ferret and mouse brain slices, rhythmic network events can be generated by perfusion with ACSF containing a low Ca^2+^/high K^+^ concentration [24, 49-51]. Here, we observed GDPs in layer 2/3 of human neocortex perfused with a standard, widely used ACSF suggesting that human neocortex is more prone to generate synchronized network activity than rat neocortex under the same conditions. In the present study, the likelihood of observing GDPs did not correlate with patient age, gender, neocortical subregion of the tissue or pathophysiological condition, implying that the recorded synchronous network events are independent and probably not pathogenic in nature. Because human brain tissue samples can be obtained only from patients requiring surgery, we acquired spared human cortex samples generated during the surgical approach of patients undergoing operations due to epilepsy or brain tumors. The cases were carefully selected, so that the surgical approach tissue was considered normal according to anatomical, macroscopic, imaging, and neuropathological criteria. Neurons included in our dataset neither showed abnormalities in firing patterns or in other electrophysiological nor in morphological properties when compared to previous description of human cortical neurons [52-55]. GDPs were only observed within a short time window (6 h) after slice preparation. In this study, we were able to begin to prepare neocortical slices within 10 min after human tissue resection. This is likely to contribute to the maintenance of the rhythmic network activity in acute human brain slices studied here.

Here, we described for the first time that GDPs were frequently observed in a specific morphological type of interneurons, namely large basket cells. Large basket cells showing GDPs display more frequent and larger normal EPSPs, resulting in a greater membrane depolarization and in turn a higher probability that the neuron reaches the threshold for AP firing. However, these interneurons show a substantially lower input resistance compared to those without GDP activity, indicating that they are less excitable and require a higher current inflow to generate APs. It is therefore likely that the synchronous network events, GDPs, are critical for eliciting APs in these large basket cells and in turn trigger feedforward inhibition of the connected neurons. Previous studies showed that rat PV-positive basket cells have a low input resistance [41, 56, 57]. By providing perisomatic inhibition onto pyramidal cells, PV basket cells have been implicated in the generation of gamma oscillations (30-120 Hz) through feedforward and feedback inhibition [1, 58]. Apart from this, inhibition of PV-positive interneurons disrupts learning-induced augmentation of delta (0.5-4 Hz) and theta oscillations (4-10 Hz) in mouse hippocampus [59]. During slow oscillations (< 1 Hz), PV-interneurons were the most active interneuron subtype during cortical ‘Up’ states in rodent neocortex [50]. GDPs observed in this study are similar to sleep-associated slow oscillations, which are primarily generated within cortical networks and are known to support memory consolidation and maintain higher frequency oscillations [60-62].

Apart from PV-positive chandelier cells, many PV-expressing interneurons show ‘basket cell’-like morphology [38]. We found that GDP-positive interneurons not only comprise PV-positive FS interneurons but also nFS interneurons which do not express PV. These interneurons may be CCK-positive interneurons displaying nFS firing pattern. While targeting the same domains of pyramidal cells as PV-expressing interneurons, CCK- and PV-positive interneurons play complementary roles in network oscillations [63, 64]. In mouse prefrontal and temporal cortex, PV- and CCK-expressing neurons constitute approximately 30-40% of all GABAergic interneurons, a percentage similar to the GDP-positive neurons we recorded among all the interneurons [65, 66]. Interneurons showing GDPs have significantly broader and longer dendrites and axons than GDP-negative interneurons. The broad and dense axonal arborizations of PV- and CCK-basket cells make them a powerful inhibitory force in human L2/3 and may therefore play a dominant role during synchronized network oscillations.

The initiation of ‘Up’ states in slow oscillations is driven by a summation of near-synchronous unitary EPSPs. With higher EPSP amplitude and/or more synapses targeting a certain postsynaptic cell, it comes more likely that EPSPs summate to a critical level to initiate an ‘Up’ state [46]. Our results support this idea since neurons showing GDPs display a larger EPSP amplitude and higher frequency of spontaneous unitary EPSPs. This may be one of the reasons why GDPs were not detected in rat neocortex since human pyramidal cells establish stronger and more reliable synaptic connections in the local circuits [31]. Recent studies revealed that human cortex has a much higher fraction of interneurons (approximately 2.5-fold) than rodent neocortex [67]. Considering GDPs are more prominently observed in interneurons rather than PCs, human neocortex is therefore more prone to generate synchronous network activity than rodent neocortex under the same conditions. Conflicting results with respect to the modulation of slow oscillations by NE were reported in previous studies [14, 15, 68-71]. Because NE and ACh have both been implicated in the promotion of wakefulness and arousal, NE is often used together with ACh to terminate slow oscillations. However, an in vivo study showed that the release of NE promotes Down-to-Up state transition, thereby inducing slow oscillations [16]. In the current study we demonstrated for the first time that NE can initiate or increase the frequency of GDPs. Several potential mechanisms may be the basis for this adrenergic regulation of GDPs. It is conceivable that NE increases the EPSP amplitude in L2/3 large basket cells of human neocortex which in turn may promote the emergence of GDPs. NE might also induce persistent firing in L2/3 of rat prefrontal cortex via activation of α_2_ receptor-linked HCN channels, which could contribute in the synchronization of neuronal activity as well [72]. However, we found that the NE-induced enhancement of GDP frequency is mediated by β-adrenergic receptor activation. It has been reported that the activation of muscarinic ACh receptors by stimulating the brainstem cholinergic nucleus abolishes slow oscillations [14, 44]. We suggest that through activating Gs protein-coupled β-adrenergic receptors, NE regulates GDPs in a way opposite to ACh via modulating Ca^2+^ and K^+^ conductance. In accordance with this hypothesis, we found that ACh can block either spontaneous or NE-induced GDPs via activating Gi protein-coupled M_4_Rs. Moreover, ACh increased the frequency of unitary EPSPs, possibly via nicotinic receptors, suggesting that ACh causes desynchronization of GDPs into unitary EPSPs and a decorrelation of responses between neurons. In summary, GDPs were promoted or suppressed by NE and ACh, respectively. The adrenergic and cholinergic modulation of GDPs drives temporal dynamics of cortical activity and controls cortical information processing and transitions between brain states. Our study provides a basis for understanding rhythmic neuronal activity in human neocortex and is of particular interest to studies investigating slow wave sleep-dependent learning and memory consolidation.

## Methods

### Patients and animals

All patients underwent neurosurgical resections because of pharmaco-resistant epilepsy or tumor removal. Written informed consent to use spare neocortical tissue acquired during the surgical approach was obtained from all patients. The study was reviewed and approved by the local ethic committee (EK067/20). For this study, we collected data from 21 patients (14 females, 7 males; age ranging from 8 to 75 years old) (Tab. 1). The cases were meticulously selected to fulfill two main criteria: 1) availability of spare tissue based on the needed surgical approach; and 2) normal appearance of the tissue according to radiological and intraoperative criteria (absence of edema, absence of necrosis, and distance to any putative intracerebral lesion). In addition, samples from tumor cases were neuropathologically reviewed to rule out the presence of tumor cells in the examined neocortical specimen.

All experimental procedures involving animals were performed in accordance with the guidelines of the Federation of European Laboratory Animal Science Association, the EU Directive 2010/63/ EU, and the German animal welfare law. In this study, Wistar rats (Charles River, either sex) aged 40–55 postnatal days were anesthetized with isoflurane and then decapitated. Rats were obtained from Charles River and kept under a 12-h light–dark cycle, with food and water available ad libitum.

### Slice preparation

Human cortex was carefully micro-dissected and resected with minimal use of bipolar forceps to ensure tissue integrity. Resected neocortical tissue from the temporal or frontal cortex was directly placed in an ice-cold artificial cerebrospinal fluid (ACSF) containing (in mM): 110 choline chloride, 26 NaHCO_3_, 10 D-glucose, 11.6 Na-ascorbate, 7 MgCl_2_, 3.1 Na-pyruvate, 2.5 KCl, 1.25 NaH_2_PO_4_, und 0.5 CaCl_2_) (325 mOsm/l, pH 7,45) and transported to the laboratory. Slice preparation commenced within 10 min after tissue resection. Pia was carefully removed from the human tissue block using forceps and the pia-white matter (WM) axis was identified. 300 µm thick slices were prepared using a Leica VT1200 vibratome in ice-cold ACSF solution containing 206 mM sucrose, 2.5 mM KCl, 1.25 mM NaH_2_PO_4_, 3mM MgCl_2_, 1 mM CaCl_2_, 25 mM NaHCO_3_, 12mM Nacetyl-L-cysteine, and 25 mM glucose (325 mOsm/l, pH 7,45). During slicing, the solution was constantly bubbled with carbogen gas (95% O_2_ and 5% CO_2_). After cutting, slices were incubated for 30 min at 31–33°C and then at room temperature in ACSF containing (in mM): 125 NaCl, 2.5 KCl, 1.25 NaH_2_PO_4_, 1 MgCl_2_, 2 CaCl_2_, 25 NaHCO_3_, 25 D-glucose, 3 myo-inositol, 2 sodium pyruvate, and 0.4 ascorbic acid (300 mOsm/l; 95% O_2_ and 5% CO_2_). To maintain adequate oxygenation and a physiological pH level, slices were kept in carbogenated ACSF (95% O_2_ and 5% CO_2_) during the transportation.

The rat brain was quickly removed and placed in an ice-cold sucrose containing ACSF. The experimental procedures used here have been described previously [73]. 300 µm thick coronal slices of the prelimbic medial prefrontal cortex (mPFC) and temporal association cortex were cut and incubated using the same procedures and solutions as described above for human slices.

### Organotypic slice cultures of human neocortex

Preparation and cultivation of slice cultures of human neocortex followed previously published protocols [74]. In brief, the neocortex was carefully micro-dissected and resected with only minimal use of bipolar forceps to ensure tissue integrity, directly transferred into ice-cold artificial cerebrospinal fluid (aCSF) (in mM: 110 choline chloride, 26 NaHCO_3_, 10 D-glucose, 11.6 Na-ascorbate, 7 MgCl_2_, 3.1 Na-pyruvate, 2.5 KCl, 1.25 NaH_2_PO_4_, und 0.5 CaCl_2_) equilibrated with carbogen (95% O_2_, 5% CO_2_) and immediately transported to the laboratory. Tissue was kept always submerged in cool and carbogenated aCSF. After removal of the pia, tissue chunks were trimmed perpendicular to the cortical surface and 250 µm thick slices were prepared using a live tissue vibratome. After the cortical tissue was sliced as described above, slices were cut into several evenly sized pieces. Subsequently, slices were transferred onto culture membranes (uncoated 30 mm Millicell-CM tissue culture inserts with 0.4 µm pores, Millipore) and kept in six-well culture dishes (BD Biosciences). For the first hour following the slicing procedure, slices were cultured on 1.5 ml intermediate step HEPES media (48% DMEM/F-12 (Life Technologies), 48% Neurobasal (Life Technologies), 1x N-2 (Capricorn Scientific), 1x B-27 (Capricorn Scientific), 1x Glutamax (Life Technologies), 1x NEAA (Life Technologies) + 20 mM HEPES before changing to 1.5 ml hCSF per well without any supplements. No antibiotics or antimycotics were used during cultivation. The plates were stored in an incubator (MCO-170AICUVH-PE, PHC Corporation) at 37°C with 5% CO_2_ and 100% humidity. For MEA recordings, slice cultures were transferred into the recording chamber of a MEA Setup (described below).

### Whole-cell recordings

Whole cell recordings were performed in acute slices 30 hours at most after slice preparation for human brain tissues and within 8 hours for rat brains. During whole-cell patch-clamp recordings, human or rat slices were continuously perfused (perfusion speed # 5 ml/min) with ACSF bubbled with carbogen gas and maintained at 30–33°C. Patch pipettes were pulled from thick wall borosilicate glass capillaries and filled with an internal solution containing (in mM): 135 K-gluconate, 4 KCl, 10 HEPES, 10 phosphocreatine, 4 Mg-ATP, and 0.3 GTP (pH 7.4 with KOH, 290–300 mOsm). Neurons were visualized using either Dodt gradient contrast or infrared differential interference contrast microscopy. Human layer 2/3 neurons were identified and patched according to their somatic location (300–1200 µm from pia) [74]. In rat acute prelimbic cortical slices, layer 2 is clearly distinguishable as a thin dark band that is densely packed with neuron somata. Layer 3 is about 2–3 times wider than layer 2 and has about the same width as layer 1. According to previous publications, layer 2/3 was located at a depth of 200 to 550 µm from the pia [75]. Putative PCs and interneurons were differentiated on the basis of their intrinsic action potential (AP) firing pattern during recording and after post hoc histological staining also by their morphological appearance.

Whole-cell patch clamp recordings of human or rat layer 2/3 neurons were made using an EPC10 amplifier (HEKA). Signals were sampled at 10 kHz, filtered at 2.9 kHz using Patchmaster software (HEKA), and later analyzed offline using Igor Pro software (Wavemetrics). Recordings were performed using patch pipettes of resistance between 5 and 10 MΩ. Biocytin was added to the internal solution at a concentration of 3–5 mg/ml to stain patched neurons. A recording time >15 min was necessary for an adequate diffusion of biocytin into dendrites and axons of patched cells [76].

### Multi-electrode array (MEA) recordings

To perform the MEA recordings of the human cortical cultures, the brain slice was excised from the insert with the slice still attached to the culturing membrane. Subsequently, the slice was moved to the MEA chamber and placed onto the electrodes of the MEA Chip with the slice surface facing down. For fixation and improved contact with the electrodes, the slice was fixed in place by a weighted, close-meshed harp (ALA-HSG MEA-5BD, Multi Channel Systems MCS GmbH). Slices equilibrated at least 30 min on the chip with constant carbogenated ACSF (same as used for acute slices) perfusion at 30-33°C before MEA recordings were started. MEA recordings were performed using a 256–MEA (16 × 16 lattice) with electrode diameter of 30 µm and electrode spacing of 200 µm, thus covering a recording area of #3.2 × 3.2 mm^2^ (USB-MEA 256-System, Multi Channel Systems MCS GmbH). Recordings with the 256-MEA were performed at a sampling rate of 10-25 kHz using the Multi-Channel Experimenter (Multi Channel Systems MCS GmbH).

### Drug application

NE (10 µM or 30 µM) and ACh (10 µM, 15 µM or 30 µM) were bath applied for 150–300 s through the perfusion system during whole-cell patch clamp or MEA recordings. In a subset of human neurons, propranolol (20 µM), tropicamide (TRO, 1 µM), tetrodotoxin (TTX, 0.5 µM), cyanquixaline (CNQX, 10µM), mecamylamine (1 µM) or prazosin (2 µM) were bath applied for 200–600 s to study the pharmacological mechanisms. Drugs were purchased from Sigma-Aldrich or Tocris.

### Histological staining

After recordings, brain slices containing biocytin-filled neurons were fixed for at least 24 h at 4°C in 100 mM phosphate buffer solution (PBS, pH 7.4) containing 4% paraformaldehyde (PFA). After rinsing several times in 100 mM PBS, slices were treated with 1% H_2_O_2_ in PBS for about 20 min to reduce any endogenous peroxidase activity. Slices were rinsed repeatedly with PBS and then incubated in 1% avidin-biotinylated horseradish peroxidase (Vector ABC staining kit, Vector Lab. Inc.) containing 0.1% Triton X-100 for 1 h at room temperature. The reaction was catalyzed using 0.5 mg/ml 3,3-diaminobenzidine (DAB; Sigma-Aldrich) as a chromogen. Subsequently, slices were rinsed with 100 mM PBS, followed by slow dehydration with ethanol in increasing concentrations, and finally in xylene for 2–4 h. After that, slices were embedded using Eukitt medium (Otto Kindler GmbH).

In a subset of experiments, we tried to identify the expression of the molecular marker - a calcium-binding protein parvalbumin (PV) in human layer 2/3 interneurons. To this end, during electrophysiological recordings, Alexa Fluor 594 dye (1:500, Invitrogen) was added to the internal solution for post hoc identification of patched neurons. After recording, slices (300 µm) were fixed with 4% PFA in 100 mM PBS for at least 24 h at 4°C and then permeabilized in 1% milk powder solution containing 0.5% Triton X-100 and 100 mM PBS. Primary and secondary antibodies were diluted in the permeabilization solution (0.5% Triton X-100 and 100 mM PBS) shortly before the antibody incubation. For single-cell PV staining, slices were incubated overnight with Rabbit-anti-PV primary antibody (1:120, ab11427, Abcam) at 4°C and then rinsed thoroughly with 100 mM PBS. Subsequently, slices were treated with Donkey-anti-Rabbit Alexa Fluor secondary antibodies (1:400, A21207, Invitrogen) for 2–3 h at room temperature in the dark. After rinsing with 100 mM PBS, the slices were embedded in Fluoromount. Fluorescence images were taken using the Olympus CellSens platform. The position of the patched neurons was identified by the biocytin conjugated Alexa dye so that the expression of PV could be examined in biocytin-stained neurons. After acquiring fluorescent images, slices were incubated in 100 mM PBS overnight and then used for subsequent histological processing as described above.

### Morphological 3D reconstructions

Using NEUROLUCIDA® software (MBF Bioscience, Williston, VT, USA), morphological reconstructions of biocytin filled human layer 2/3 interneurons were made at a magnification of 1000-fold (100-fold oil-immersion objective and 10-fold eyepiece) on an upright microscope. Neurons were selected for reconstruction based on the quality of biocytin labelling when background staining was minimal. Neurons with major truncations due to slicing were excluded. Embedding using Eukitt medium reduced fading of cytoarchitectonic features and enhanced contrast between layers [76]. This allowed the reconstruction of different layer borders along with the neuronal reconstructions. Furthermore, the position of soma and layers were confirmed by superimposing the Dodt gradient contrast or differential interference contrast images taken during the recording. The tissue shrinkage was corrected using correction factors of 1.1 in the x–y direction and 2.1 in the z direction [76]. Analysis of 3D-reconstructed neurons was done with NEUROEXPLORER® software (MBF Bioscience, Williston, VT, USA).

### Data analysis

#### Single-cell recording data analysis

Custom written macros for Igor Pro 6 (WaveMetrics) were used to analyze the recorded electrophysiological signals. The resting membrane potential (V_m_) of the neuron was measured directly after breakthrough to establish the whole-cell configuration with no current injection. The input resistance was calculated as the slope of the linear fit to the current–voltage relationship. For the analysis of single spike characteristics such as threshold, amplitude and half-width, a step size increment of 10 pA for current injection was applied to ensure that the AP was elicited very close to its rheobase current. The spike threshold was defined as the point of maximal acceleration of the membrane potential using the second derivative (d2V/dt2), which is, the time point with the fastest voltage change. The spike amplitude was calculated as the difference in voltage from AP threshold to the peak during depolarization. The spike half-width was measured as the time difference between rising phase and decaying phase of the spike at half-maximum amplitude.

The spontaneous activity was analyzed using the program SpAcAn (https://www.wavemetrics.com/ project/SpAcAn). Normal EPSPs and GDPs were distinguished by dramatic differences in event amplitude and decay time. A threshold of 0.2 mV was set manually for detecting EPSP events while a threshold of 3 mV was set for detecting GDPs. Recordings were not filtered to reduce noise before data analysis. When marking normal EPSPs, small EPSPs distributing in decay phrase but not rising phrase of GDPs were included into analysis. To study oscillatory network activities, we computed time-frequency representations of the signals by performing wavelet analysis using the Time-Frequency Toolkit (https://www.wavemetrics.com/project/TFPlot). Morlet wavelets were used for decomposition of recording signals as they provide an ideal compromise between time and frequency resolution [77].

#### MEA data analysis

MEA recordings were analyzed using custom-written programs in Python, detecting and quantifying the mean firing rate, number of active channels, bursting channels, and network bursts. First, the raw signal was filtered using a band-pass filter (Butterworth 2nd order). The spike identification was performed according to a threshold-based method using median absolute deviation (MAD) / 0.6745 x -5. Signal deviations were detected and aligned to the next minimum of the signal with a 1 ms dead time. The firing rate for each recording was defined as the number of recorded spikes divided by the duration of the recording (in s). Bursting channels were calculated using a modified adaptive network-wide cumulative moving average (CMA) approach in which all the electrodes of one MEA in multiple measurement time points were analyzed together [78].

In short, all inter-spike intervals (ISIs) were calculated and grouped into 5 ms bins for the whole recording. Next, a CMA over the histogram of the bins was calculated and a burst threshold was determined, therefore adapting the detection of bursts to the basic activity of the entire slice [78]. The detection sensitivity was further increased by applying a minimum and maximum threshold of 60 ms and 140 ms, respectively. Whenever the threshold was undercut by at least three consecutive spikes, it was defined as a single-channel burst.

For a quantitative analysis of spiking synchronization, we used a graph theoretical approach to identify the degree of centrality of active channels [79-81]. Specifically, we designated MEA contacts as nodes and the shared spiking time as edges to construct a graph representation of the recorded network activity. To construct edges, we grouped spike trains for each channel into 200 ms long bins and defined two channels as connected by an edge only if they both recorded spikes are in the same bin. By using this approach, we were able to examine the functional connectivity and activity patterns in the neuronal network.

We used mean degree centrality as a measure of the connectivity of nodes in the network (Python library NetworkX). Mean degree centrality (MDC) quantifies the average number of edges connecting a node to other nodes in the network. The degree of each node was defined by the number of edges connected to that node, i.e. the number of other MEA contacts that share a spiking time with that contact within a 200-ms bin. Finally, MDC was calculated by summing the degree of all nodes and dividing it by the total number of nodes in the network, which is

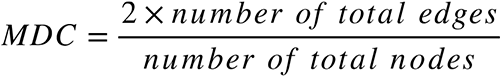

where the factor of 2 is included because each edge is counted twice (once for each node it connects). By calculating the MDC for the whole MEA recording, we were able to assess the overall connectivity of the network and identify how nodes connectivity (used as a surrogate for synchronicity) changed upon application of NE and ACh compared to baseline.

Detection of LFPs was performed after low pass filtering the signal of each channel (Butterworth 2nd order with a Nyquist frequency of 0.5 x sampling rate and a cut-off of 100 Hz); as a threshold, standard deviation of the low pass filtered signal multiplicated by three was used. Any deviation above or below this threshold with a minimum duration of 30 ms was defined as a LFP. The respective maximum and minimum deviation was defined as the amplitude of respective LFP.

All Python scripts are available on GitHub page (https://github.com/jonasort/MEA_analysis/tree/ main/modified_common_script).

#### Statistical analysis

Data was either presented as box plots (n ≥ 10) or as bar histograms (n < 10). For box plots, the interquartile range (IQR) is shown as box, the range of values within 1.5$IQR is shown as whiskers and the median is represented by a horizontal line in the box; for bar histograms, the mean ± SD is given. Wilcoxon Mann-Whitney U test was performed to access the difference between individual clusters. Statistical significance was set at P < 0.05, and n indicates the number of neurons/slices analyzed.

## Author contributions

D.F., D.Y., G.Q. and H.K. designed the experiments. D.Y. and G.Q. carried out the patch-clamp recording experiments from human and rat slices and electrophysiological data analysis. D.Y. performed Neurolucida reconstructions and performed morphological analysis. D.D performed surgeries on human patients. D.Y., A.B. and H.K. prepared acute human brain slices and A.B. prepared the human slice cultures. J.O. and V.W. performed MEA recordings. J.O., H.K. and D.D. analyzed MEA data. D.Y. and D.F. wrote the manuscript with the inputs from all authors. All authors have given approval for the final version of the manuscript.

## Competing interests

The authors declare no competing interests.

### Acknowledgement

We would like to thank Werner Hucko and Birgit Gittel for excellent technical assistance. We thank Dr. Karlijn van Aerde for custom-written macros in Igor Pro software. We are grateful for funding support from the European Union’s Horizon 2020 Framework Programme for Research and Innovation under the Framework Partnership Agreement No. 650003 (HBP FPA) to D.F., funding from DFG FOR2715, Chan Zuckerberg Initiative DAF (2020-221779) to H.K. and funding from BMBF (German Ministry of Education and Research, project number 031L0260B) to D.D..

