## Supplementary Materials for "Modulation of Giant Depolarizing Potentials (GDPs) in Human Large Basket Cells by Norepinephrine and Acetylcholine"

#### Supplementary figures and tables

Danqing Yang[1], Guanxiao Qi[1], Jonas Ort[2,7,8], Victoria Witzig[3], Aniella Bak[4], Daniel Delev[2,7,8], Henner Koch[4] and Dirk Feldmeyer [1,5,6 #]

1 Research Center Juelich, Institute of Neuroscience and Medicine 10, Research Center Juelich, 52425 Juelich, Germany.
2 Department of Neurosurgery, Faculty of Medicine, RWTH Aachen University Hospital, Aachen, Germany
3 Department of Neurology, RWTH Aachen University Hospital, 52074 Aachen, Germany.
4 Department of Neurology, Section Epileptology, RWTH Aachen University Hospital, 52074 Aachen, Germany.
5 Department of Psychiatry, Psychotherapy, and Psychosomatics, RWTH Aachen University Hospital, 52074 Aachen, Germany.
6 Jülich-Aachen Research Alliance, Translational Brain Medicine (JARA Brain), Aachen, Germany.
7 Neurosurgical Artificial Intelligence Laboratory Aachen (NAILA), RWTH Aachen University Hospital, 52074 Aachen, Germany
8 Center for Integrated Oncology, Universities Aachen, Bonn, Cologne, Düsseldorf (CIO ABCD), Germany.

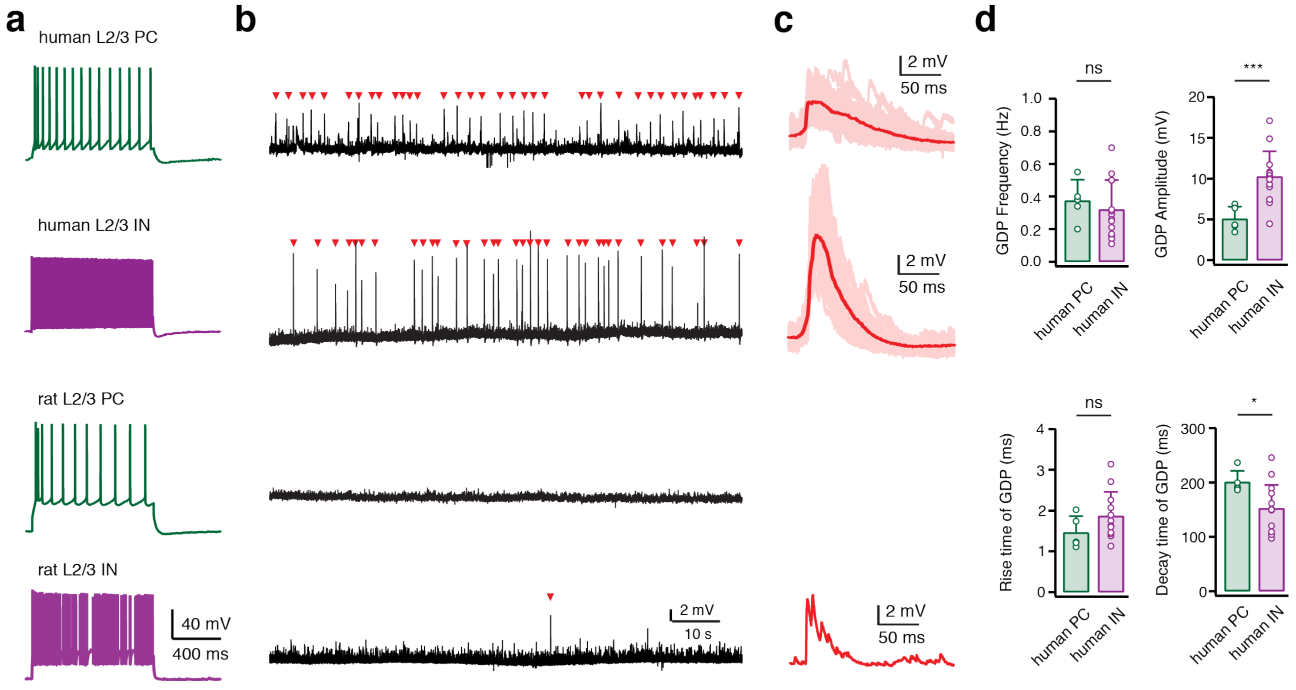


**Fig. S1 Regular giant network events were not observed in L2/3 neurons of adult rat prefrontal or temporal cortex.**

**a** Representative firing patterns of a human L2/3 PC, a human L2/3 interneuron, a rat L2/3 PC and a rat L2/3 interneuron. Firing patterns of PCs are shown in green while those of interneurons are shown in purple.

**b** Corresponding current-clamp recording traces are obtained from the same neuron in a. Spontaneously occurring GDPs are marked in red.

**c** Average and individual GDPs are superimposed and given in a dark and light red, respectively.

**d** Bar histograms comparing frequency, amplitude, rise time and decay time of GDPs between PCs (n = 5) and interneurons (n = 14). * $P < 0.05$, *** $P < 0.001$ for the Wilcoxon Mann–Whitney U test.

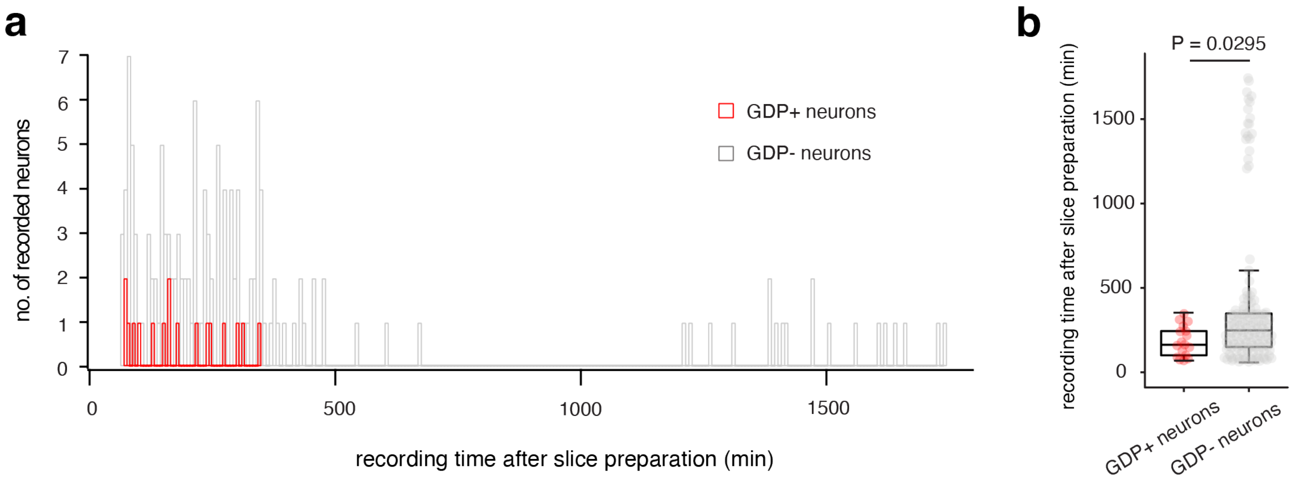


**Fig. S2 GDPs were observed only within 6 hours after slice preparation**

**a** Recording time after slice preparation for human L2/3 neurons with (red) and without (gray) GDP activity. Histograms were constructed with 6 min bins.

**b** Box plot comparing the recording time after slice preparation for GDP+ (n = 17) and GDP- (n = 146) neurons. P =0.0295 for the Wilcoxon Mann–Whitney U test.

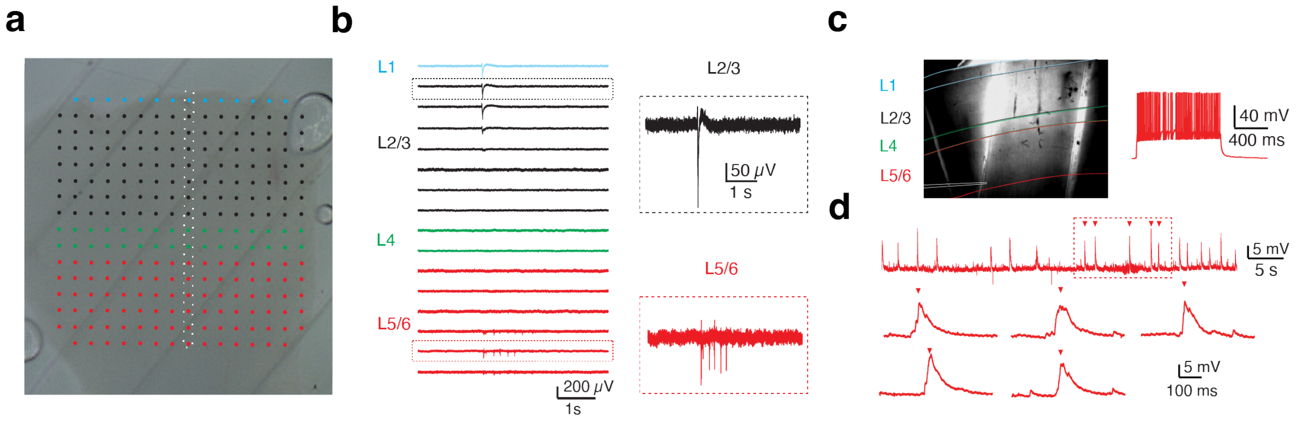


**Fig. S3 Synchronous network events were observed in layer 6.**

**a** A typical human slice culture is shown with all cortical layers (L1–L6) and overlay of the MEA on the slice. The MEA channels in L1, L2/3, L4 and L5/6 are marked by light blue, black, green and red dots, respectively.

**b** Representative voltage traces from 16 MEA channels. Black and red insets show a detailed view of LFPs in L2/3 and L5/6, respectively.

**c** Position of the recorded layer 6 interneuron in a human acute slice. Layer borders are indicated by the same color code as in (b). Firing pattern of the recorded neuron is shown on the right.

**d** Whole cell intracellular recordings from a L6 interneuron in human neocortex showing GDP activity. Five marked consecutive GDPs are shown at higher magnification at the bottom.

| | GDP+ L2/3 INs | GDP- L2/3 INs | Mann-Whitney Test |
|---|---|---|---|
| | **Morphological properties (n = 8 for each group)** | | |
| **somatic area (µm²)** | 170.2 ± 44.1 | 186.2 ± 84.1 | 0.6454 |
| **dendritic length (µm)** | 4171.2 ± 1210.4 | 2080.1 ± 1105.0 | ****0.0207*** |
| **axonal length (µm)** | 40778.6 ± 8197.8 | 12485.7 ± 5402.8 | ******0.0003*** |
| **no. of dendrites** | 6.0 ± 1.6 | 3.9 ± 1.2 | 0.3915 |
| **H-fieldspan of dendrite (µm)** | 546.2 ± 176.1 | 212.1 ± 104.0 | ******0.0003*** |
| **V-fieldspan of dendrite (µm)** | 652.4 ± 193.8 | 308.3 ± 216.8 | *****0.0047*** |
| **H-fieldspan of axon(µm)** | 1227.6 ± 427.3 | 604.0 ± 405.0 | ****0.0140*** |
| **V-fieldspan of axon (µm)** | 1007.5 ± 284.1 | 859.3 ± 404.7 | 0.4634 |
| | **Electrophysiological properties (n = 13 for each group)** | | |
| **resting membrane potential (mV)** | -67.0 ± 6.4 | -66.0 ± 6.8 | 0.7399 |
| **Input resistance (MΩ)** | 192.4 ± 85.1 | 298.5 ± 117.9 | ****0.0278*** |
| **AP half- width (ms)** | 0.48 ± 0.21 | 0.37 ± 0.12 | 0.2406 |
| **AP amplitude (mV)** | 79.2 ± 9.5 | 84.4 ± 7.5 | 0.3338 |
| **AP threshold (mV)** | -41.2 ± 2.9 | -42.3 ± 3.2 | 0.3760 |
| **AHP amplitude (mV)** | 23.7 ± 5.1 | 25.0 ± 3.7 | 0.7689 |
| **AP latency (ms)** | 196.8 ± 155.5 | 143.6 ± 101.1 | 0.7283 |
| **frequency- current slope (Hz/100 pA)** | 30.2 ± 8.7 | 40.7 ± 17.6 | 0.1164 |
| **Adaptation ($ISI_2/ISI_{10}$)** | 0.75 ± 0.24 | 0.79 ± 0.24 | 0.6981 |

**Tab. S1 Morphological and electrophysiological properties of GDP+ and GDP- human L2/3 interneurons.**

Italic bold font indicates significant differences; *P < 0.05, **P < 0.01, ***P < 0.001 for Wilcoxon Mann-Whitney U test.

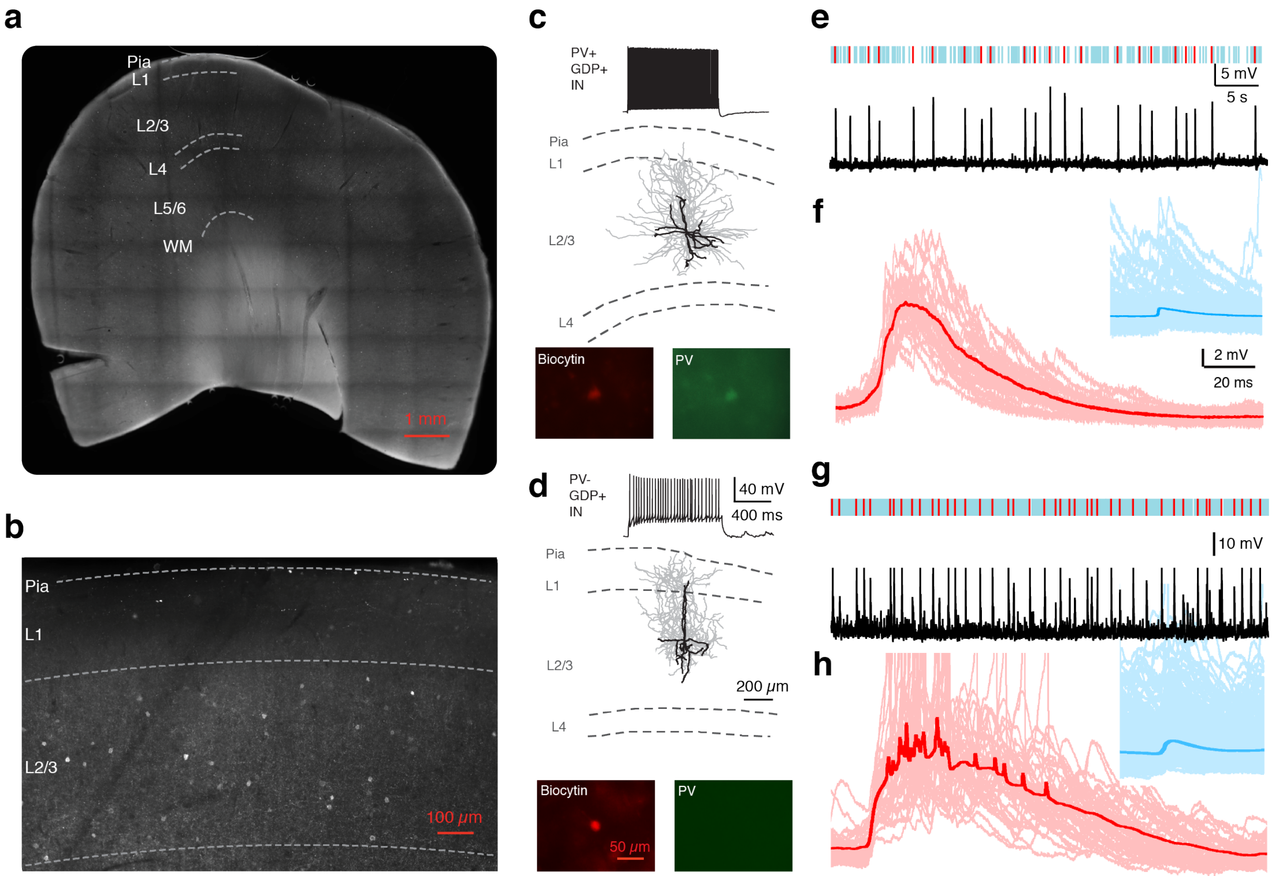


**Fig. S4 Large basket cells showing GDPs in L2/3 human cortex can be either parvalbumin (PV) positive or negative.**

**a** Representative example of a human brain slice after antibody labelling for PV. Layer borders are indicated by dashed white lines.

**b** PV positive interneurons are more abundant in layer 2/3 than in layer 1. Layer borders are indicated with dashed white lines.

**c, d** Top: Representative firing patterns of a PV+, GDP+ human L2/3 interneuron (c) and a PV- GDP+ human L2/3 interneuron (d). Middle: Corresponding morphological reconstructions of the same neurons shown above. The somatodendritic domain is given in black and the axons in gray. Bottom: Neurons were recorded using whole-cell patch-clamp technique and simultaneously filled with biocytin and the fluorescent dye Alexa 594 (red). Antibody labeling was performed to test for the expression of PV (green).

**e, g** A 50 s recording was obtained from the same PV+ (e) and PV- interneuron (g), respectively. Normal EPSPs are marked in blue while GDPs are marked in red.

**f, h** Left: The average and individual GDPs are superimposed and given in a darker and lighter red, respectively. Right: The average and individual EPSPs are superimposed and given in a

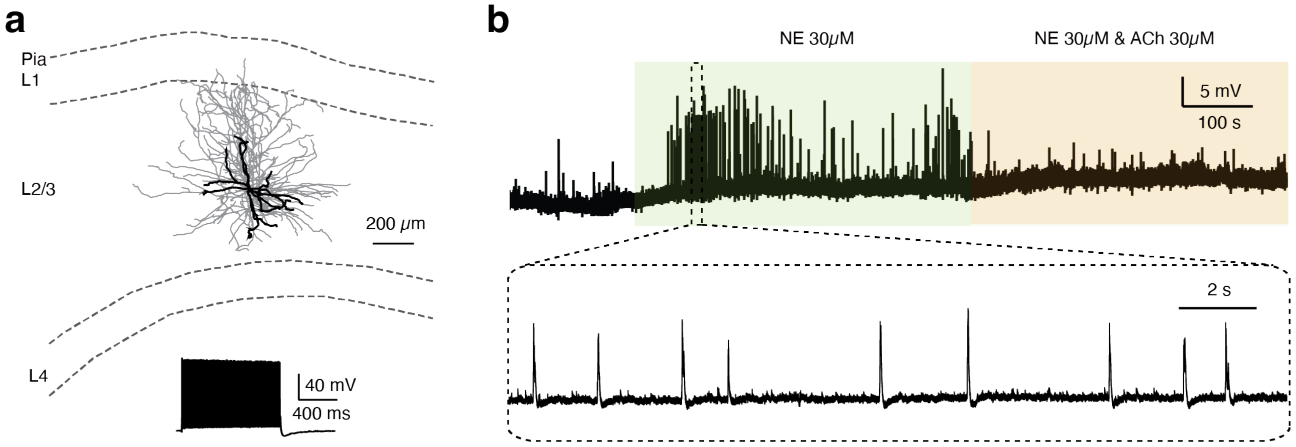


**Fig. S5 ACh suppresses the NE-induced GDPs in L2/3 human interneurons**

**a** Top: Representative morphological reconstruction of a GDP+ human L2/3 interneuron. Bottom: Corresponding firing pattern of the same neuron shown above.

**b** Representative current-clamp recordings with bath application of 30 μM NE causing an increase in GDP frequency in the same neuron shown in a. The NE effect is reversed by 30 μM ACh.

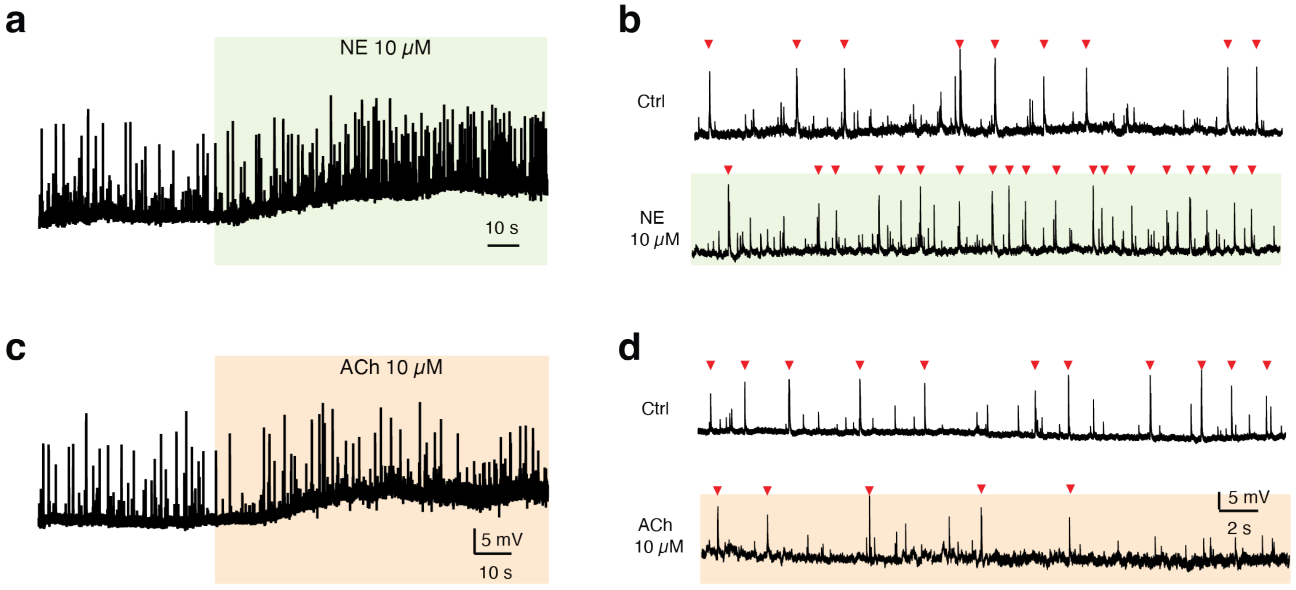


**Fig. S6 Low concentrations ACh and NE (10μM) show similar modulation of GDPs in L2/3 human interneurons**

**a** Representative current-clamp recordings with bath application of 10 μM NE showing an increase of GDP frequency and a membrane potential depolarization in a human L2/3 interneuron.

**b** Enlarged recording traces under control and 10 μM NE conditions. GDPs are marked by red arrowheads.

**c** Representative current-clamp recordings with bath application of 10 μM ACh showing a decrease of GDP frequency and a membrane potential depolarization in a human L2/3 interneuron.

**d** Enlarged recording traces under control and 10 μM ACh conditions. GDPs are marked by red arrowheads.

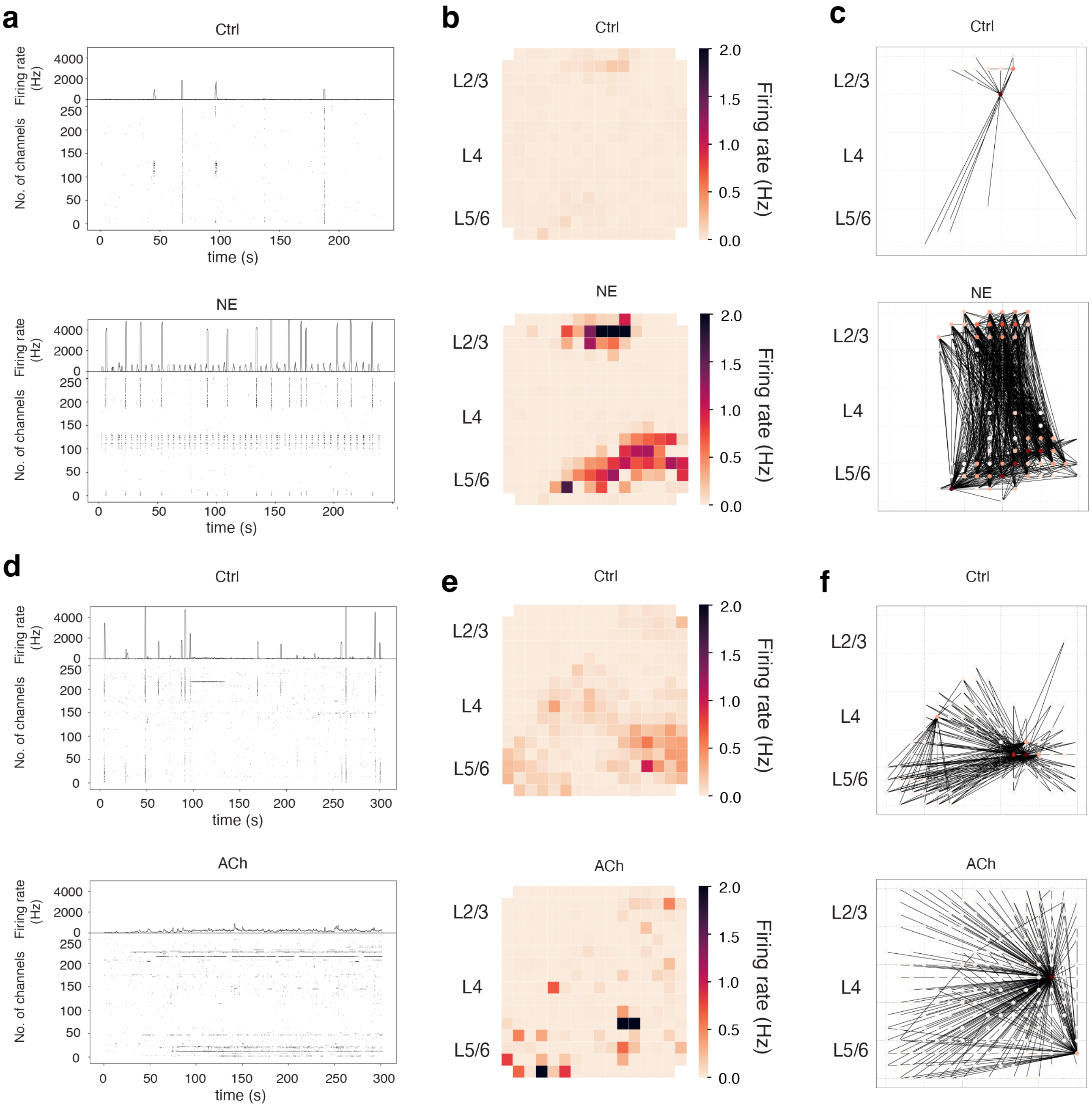


**Fig. S7. NE causes synchronization while ACh promotes desynchronization of neuronal firing.**

**a** Raster plots of the detected action potentials over a 5 min recording period under control condition and in the presence of NE showing an increase in synchronous firing.

**b** Heatmap of the average firing rate over the MEA grid from a 5 min recording in control condition and in the presence of NE showing an increase synchronous firing in L2/3 and deep layers.

**c** Graph analysis of MEA recordings over a 5 min period under control condition and in the presence of NE showing an increase of degree of centrality. A line connecting two MEA channels (*nodes of the graph*) represents an *edge of the graph*. Two *nodes* are connected by one *edge* only if they spike synchronously within a bin of 200 ms. Degree centrality is calculated from number of connected *nodes* and number of *edges* used for these connections.

**d** Raster plots of the detected action potentials over the 5 min recording period under control condition and in the presence of ACh showing a decrease in synchronous firing.

**e** Heatmap of the average firing rate over the MEA grid from a 5 min recording in control condition and in the presence of ACh showing a decrease in temporal and spatial correlation of global AP firing.

**f** Graph analysis of MEA recordings over a 5 min period under control condition and in the presence of ACh showing a decrease of degree of centrality. For details s. c).

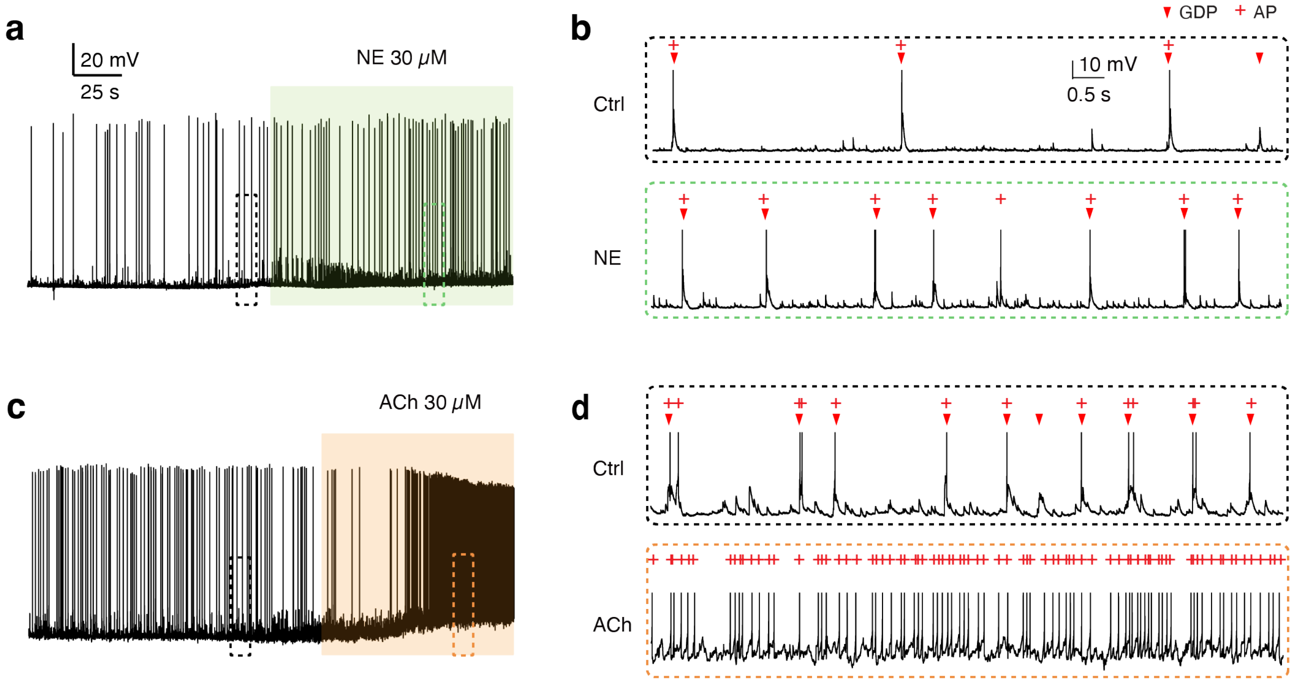


**Fig. S8. NE and ACh increase AP firing rates via different mechanisms**

**a** Representative current-clamp recordings with bath application of 30 μM NE showing an increase of GDP frequency and AP firing rates in a human L2/3 interneuron.

**b** Enlarged recording traces under control and 30 μM NE conditions. GDPs and APs are marked by red arrowheads and red plus mark, respectively.

**c** Representative current-clamp recordings with bath application of 30 μM ACh showing a strong membrane potential depolarization and increase of AP firing rates in a human L2/3 interneuron.

**d** Enlarged recording traces under control and 30 μM ACh conditions. GDPs and APs are marked by red arrowheads and red plus mark, respectively.
